# Local human movement patterns and land use impact exposure to zoonotic malaria in Malaysian Borneo

**DOI:** 10.1101/734590

**Authors:** Kimberly M Fornace, Neal Alexander, Tommy R Abidin, Paddy M Brock, Tock H Chua, Indra Vythilingam, Heather M. Ferguson, Benny O. Manin, Meng L. Wong, Sui Hann Ng, Jon Cox, Chris J Drakeley

**Affiliations:** Faculty of Infectious and Tropical Diseases, London School of Hygiene and Tropical Medicine, London, UK; Department of Infectious Disease Epidemiology, London School of Hygiene and Tropical Medicine, London, UK; Department of Pathobiology and Medical Diagnostics, Faculty of Medicine and Health Sciences, Universiti Malaysia Sabah, Kota Kinabalu, Malaysia; Institute of Biodiversity, Animal Health and Comparative Medicine, College of Medical, Veterinary and Life Sciences, University of Glasgow, UK; Parasitology Department, Faculty of Medicine, University of Malaya, Kuala Lumpur, Malaysia

**Keywords:** Disease ecology, spatial epidemiology, *Plasmodium knowlesi*, malaria, human movement, land use

## Abstract

Human movement into insect vector and wildlife reservoir habitats determines zoonotic disease risks; however, few data are available to quantify the impact of land use on pathogen transmission. Here, we utilise GPS tracking devices and novel applications of ecological methods to develop fine-scale models of human space use relative to land cover to assess exposure to the zoonotic malaria *Plasmodium knowlesi* in Malaysian Borneo. Combining data with spatially explicit models of mosquito biting rates, we demonstrate the role of individual heterogeneities in local space use in disease exposure. At a community level, our data indicate that areas close to both secondary forest and houses have the highest probability of human *P. knowlesi* exposure, providing quantitative evidence for the importance of ecotones. Despite higher biting rates in forests, incorporating human movement space use into exposure estimates illustrates the importance of intensified interactions between pathogens, insect vectors and people around habitat edges.

## Introduction

Environmental change and human encroachment into wildlife habitats are key drivers in the emergence and transmission of zoonotic diseases (1, 2). Individual movements into different habitats influence exposure to disease vectors and animal reservoirs, determining risk and propagation of vector-borne diseases (3–5). Increased contact between these populations is theorised to drive increases of the zoonotic malaria *Plasmodium knowlesi* in Malaysian Borneo, now the main cause of human malaria within this region. *P. knowlesi* is carried by long- and pig-tailed macaques (*Macaca fascicularis* and *M. nemestrina*) and transmitted by the *Anopheles leucospryphus* mosquito group, both populations highly sensitive to land cover and land use change (6). Although higher spatial overlap between people, macaques and mosquito vectors likely drives transmission, the impact of human movement and land use in determining individual infection risks is poorly understood (7).

The emergence of the zoonotic malaria *Plasmodium knowlesi* has been positively associated with both forest cover and historical deforestation (8, 9). However, out of necessity, statistical approaches to assess environmental risk factors for *P. knowlesi* and other infectious diseases typically evaluate relationships between disease metrics and local land cover surrounding houses or villages. While an individual may spend most of their time within the vicinity of their residence, this area does not necessarily represent where they are most likely to be exposed to a disease. This is supported by varying associations between *P. knowlesi* occurrence and landscape variables at different distances from households, ranging from 100m to 5km, likely partially due to human movement into different surrounding habitats (8, 10). Although land cover variables describing physical terrestrial surfaces are frequently incorporated into disease models, land use is rarely quantified. Land use is commonly defined as “the arrangements, activities, and inputs that people undertake in certain land cover types” (11). Places with the similar types of land cover may be used very differently, with the activities and frequencies with which people visit these places determining the spatial distribution of disease (1).

Mathematical modelling studies have revealed the importance of spatial variation in contact rates due to the movement of individuals through heterogeneous environments with varying transmission intensity (12). A multi-species transmission model of *P. knowlesi* highlighted the role of mixing patterns between populations in different ecological settings in determining the basic reproductive rate and subsequent modelling studies illustrate the sensitivity of this disease system to population densities of both people and wildlife hosts (7, 13). However, although mechanistic models have been extended to explore the potential importance of these heterogeneities in disease dynamics, there are inherent constraints on model complexity and most models make simplistic assumptions about the habitat uses of different populations.

Empirical data on human population movement is increasingly available, allowing assessment of the impact of mobility on infectious disease dispersion and risks (5). On larger spatial scales, mobile phone data has revealed the role of human migration in the transmission of infectious diseases such as malaria, dengue and rubella (14–16).Although this data can provide insights into long range movements, spatial resolution of this data is limited, particularly in areas with poor or no mobile coverage, such as forested areas (17). Alternatively, the advent of low-cost GPS tracking devices allows quantification of fine-scale movements, demonstrating marked heterogeneity in individual movement and risk behaviours (3, 18). Combining these data with detailed data on land cover and vector dynamics can provide new insights into how landscapes affect *P. knowlesi* transmission.

Previous studies of *P. knowlesi* have relied on questionnaire surveys, identifying self-reported travel to nearby plantations and forest areas as a risk factor for *P. knowlesi* and other malaria infections (e.g. (19–21)). However, the resultant spatial range and frequency of these movements remain unknown and the definition of different habitat types is entirely subjective. Further, little is known about differences in local movement patterns in different demographic groups. While infections in male adults have been linked to forest and plantation work, it is unknown whether infections reported in women and young children are likely to arise from exposure to similar environments (22). The main mosquito vector in this area, *An. balabacensis*, is primarily exophagic and has been identified in farm, forest and village areas near houses (23, 24). Macaque populations are reported in close proximity to human settlements and molecular and modelling studies suggest transmission remains primarily zoonotic in this area (7, 25, 26). A case control study detected higher abundances of *An. balabacensis* near *P. knowlesi* case housesholds, suggesting the possibility of peri-domestic transmission (24). Understanding the importance of these habitats is essential to effectively target intervention strategies and predict impacts of future environmental changes.

Key questions remain about where individuals are likely to be exposed to *P. knowlesi* and how landscape determines risk. Functional ecology approaches allow the distribution of different populations to be modelled based on biological resources and relate transmission to landscape and environmental factors (27). Within wildlife ecology, numerous methods have been developed to estimate utilisation distributions (UDs), the probability of an individual or species being within a specific location during the sampling period (28). Although these methods traditionally rely on kernel density smoothing, kernel density estimates may not actually reflect time individuals spend in a specific location if there is substantial missing data or irregular time intervals. Alternatively, biased random bridges (BRBs) improve on these methods by estimating the UD as a time-ordered series of points, taking advantage of the autocorrelated nature of GPS tracks to bias movement predictions towards subsequent locations in a time series (29). This allows for interpolation of missing values and adjustment for spatial error to estimate UDs representing both the intensity (mean residence time per visit) and frequency of individual visits to specific locations. By integrating these estimates of individual space use with detailed spatial and environmental data in a Bayesian framework, fine-scale patterns of human land use can be predicted and overlaid with spatiotemporal models of mosquito distribution. This allows exploration how landscape composition, as well as configuration and connectivity between habitats, impacts human exposure to *P. knowlesi* and other vector-borne and zoonotic diseases.

Focusing on one aspect of land use, human movement and time spent within different land cover types, we explored the role of heterogeneity in local space use on disease exposure. Rolling cross-sectional GPS tracking surveys were conducted in two study areas with on-going *P. knowlesi* transmission in Northern Sabah, Malaysia (Matunggong and Limbuak (30)). We aimed to characterise local movement patterns and identify individuals and locations associated with increased *P. knowlesi* exposure risks by: 1. analysing individual movement patterns and developing predictive maps of human space use relative to spatial and environmental factors, 2. modelling biting rates of the main vector *An. balabacensis*, and 3. assessing exposure risks for *P. knowlesi* based on predicted mosquito and human densities (Figure 1) Integrating these three approaches allowed a uniquely spatially explicit examination of disease risk.

**Figure 1.**
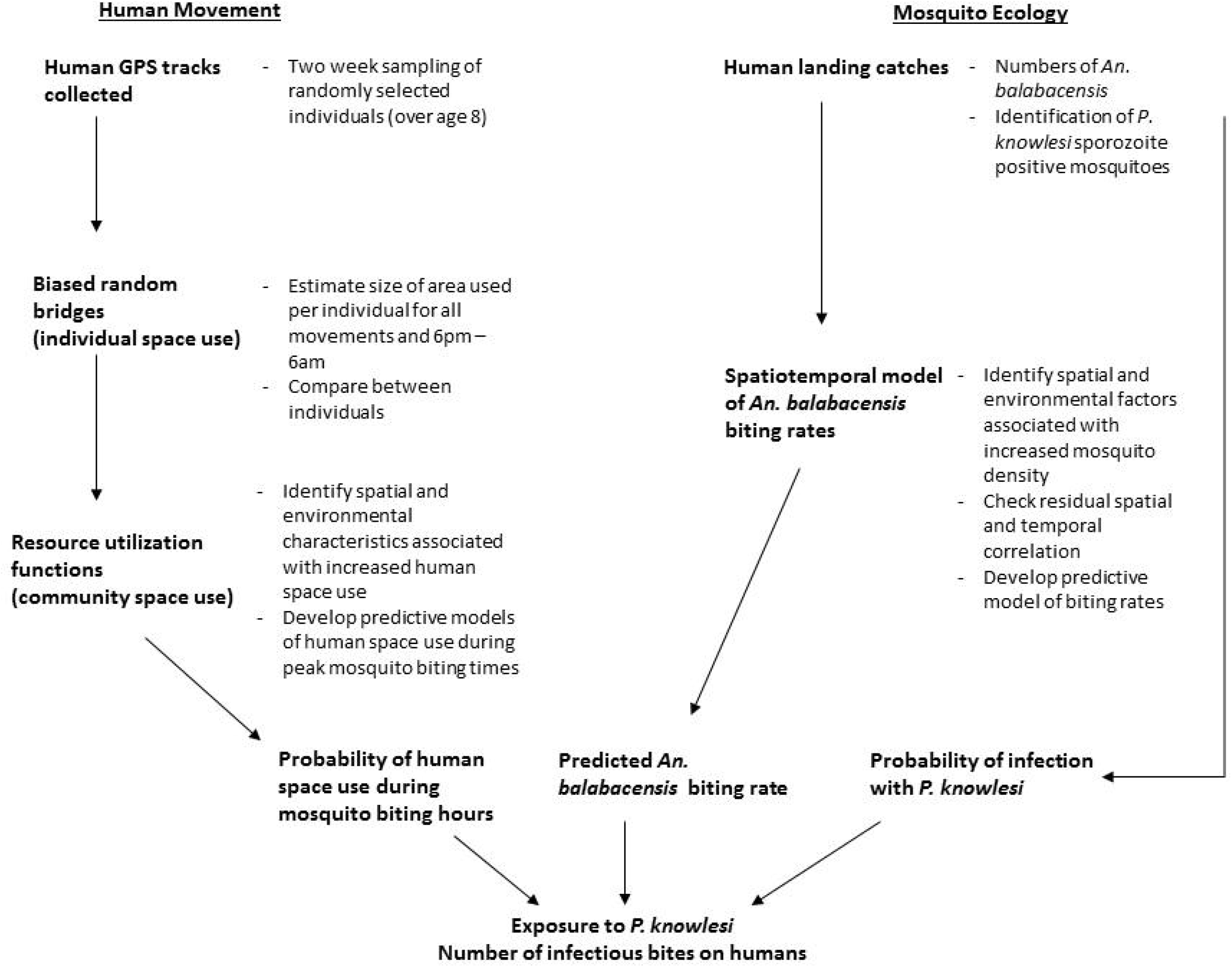
Analysis methods used to estimate individual and community-level exposure to *P. knowlesi* sporozoite positive *An. balabacencis* bites

## Methods

### Study site

This study was conducted in two rural communities in Northern Sabah, Malaysia: Matunggong, Kudat (6°47N, 116°48E, population: 1260) and Limbuak, Pulau Banggi (7°09N, 117°05E, population: 1009) (Figure 2). These areas were the focus for integrated entomology, primatology and social science studies for risk factors for *P. knowlesi* (https://www.lshtm.ac.uk/research/centres-projects-groups/monkeybar), with clinical cases and submicroscopic infections reported from both sites and *P. knowlesi* sero-prevalence estimated as 6.8% and 11.7% in Matunggong and Limbuak respectively (30).

**Figure 2.**
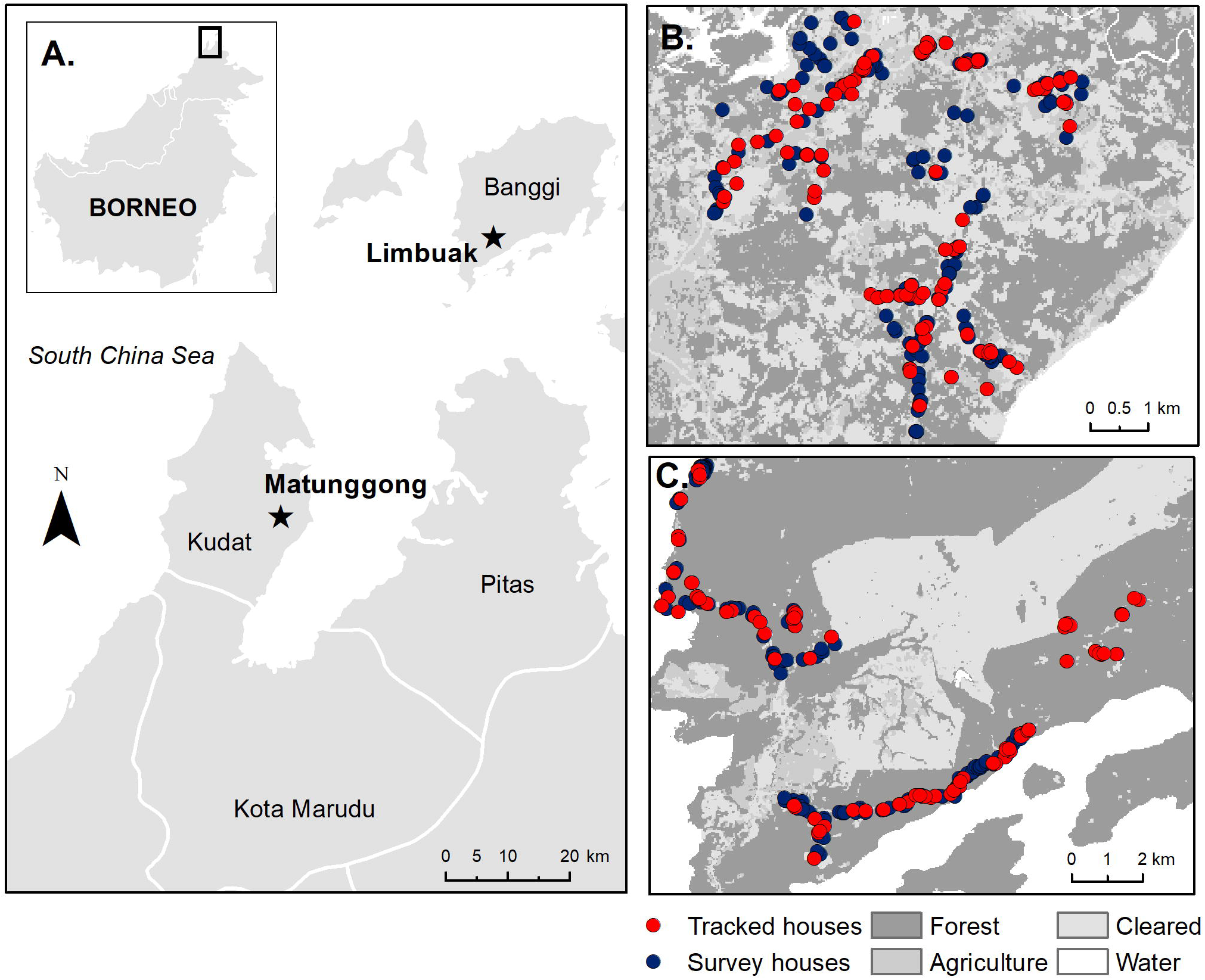
A. Location of study sites and tracked houses (households with one or more individual GPS tracked) and survey houses (households with only questionnaire data collected and used for prediction) in B. Matunggong, Kudat and C. Limbuak, Banggi; description of land cover classification and survey methodology in (30)

Demographic data and GPS locations of primary residences were collected for all individuals residing in these areas (30). Potential spatial and environmental covariates for these sites were assembled from ground-based and remote-sensing data sources (Supplementary Table 1). The enhanced vegetation index (EVI) was used to capture temporal changes in vegetation levels; this index captures photosynthetic activity and has higher sensitivity in high biomass areas compared to the normalised difference vegetation index (NDVI) frequently used. Due to the high cloud cover within this area, EVI at a high spatial resolution could not be obtained for all time periods. Instead, EVI data at a lower spatial but higher temporal resolution was used and monthly averages were calculated from all available cloud-free data and resampled to 30m per pixel (31).

### GPS tracking survey

A minimum of 50 participants per site were targeted in a rolling cross-sectional survey (32). During pre-defined two-week intervals, randomly selected participants from comprehensive lists of eligible community members were asked to carry a QStarz BT-QT13000XT GPS tracking device (QStarz, Taipei, Taiwan) programmed to record coordinates continuously at one-minute intervals for at least 14 days regardless of individual movement. Individuals were excluded if they were not primarily residing in the study area, under 8 years old or did not consent. Trained fieldworkers visited the participant every two days to confirm the device was functioning, replace batteries and administer questionnaires on locations visited and GPS use. Fieldworkers recorded whether the device was working and if the individual was observed carrying the GPS device to assess compliance. Individuals were excluded from analysis if insufficient GPS data were collected (less than 33% of sampling period) or individuals were observed not using the device for two or more visits.

### Human space use

Biased random bridges (BRBs) were used to calculate individual utilisation distributions (UDs), the probability of an individual being in a location in space within the sampled time period (29). Within this study, large proportions of GPS fixes were missed due to technical issues with batteries and GPS tracking; BRBs were used to interpolate between known locations and adjust for missing data, using the time series GPS data to provide a more accurate estimate of space use. UDs were calculated separately for each individual for all movement and night-time only movements (6pm – 6am).

To fit BRBs, we estimated the maximum threshold between points before they were considered uncorrelated (*T*_*max*_) as 3 hours based on typical reported activity times. The minimum distance between relocations (*L*_*min*_), the distance below which an individual is considered stationary, was set at 10m to account for GPS recording error based on static tests. Finally, the minimum smoothing parameter (*h*_*min*_), the minimum standard deviation in relocation uncertainty, was set as 30m to account for the resolution of habitat data and capture the range of locations an individual could occupy while being recorded at the same place (28, 29). Estimates of the core utilisation area (home range) were based on the 99^th^ percentile, representing the area with a 99% cumulative probability distribution of use by the sampled individual.

To assess relationships between space use and environmental factors and develop predictive maps of community space use, we fit resource utilisation functions, regression models in which the UDs are used as the response variable, improving on models using raw GPS count points as the response when there is location uncertainty and missing data (33). The probability density function (UD) per individual was rasterised to 30m^2^ grid cells and environmental and spatial covariates extracted for each grid cell. Potential environmental covariates included distance to the individual’s own house, distance to closest house, distance to roads, land use class (forest, agriculture, cleared or water), distance to forest edge, elevation and slope (Supplementary table 1). Resource utilisation was modelled as a Bayesian semi-continuous (hurdle) model with two functionally independent components, a Bernoulli distribution for probability of individual i visiting a specific grid cell *j* (*ω*_*ij*_) and a gamma distribution for the UD in grid cells visited (*y*_*ij*_) (34, 35). For each individual, we defined absences to be all grid cells with a UD less than 0.00001, indicating a very low probability the individual visited this grid cell during the study period. We included all presences (grid cells with a UD > 0.00001) and randomly subsampled equal numbers of absences (grid cells not visited) for each individual as including equal numbers of presences and absences can improve predictive abilities of species distribution models (36). The utilisation distribution for grid cells visited is defined as:

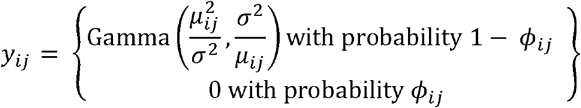

Where the mean of *y*_*ij*_ is given by:

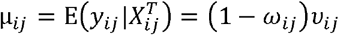

The full model was specified as:

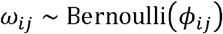

With the linear predictor for the Bernoulli model specified as:

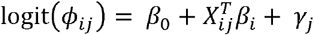

Where *β*_0_ represents the intercept, 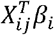 represents a vector of covariate effects and *γ*_*j*_ represents the additive terms of random effects for individual. For the Gamma component *σ*^2^ is the variance and the linear predictor *v*_*ij*_ is specified as:

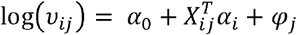

With *α*_0_ representing the intercept, 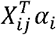 representing a vector of coordinates and *ϕ*_*j*_ representing the random effects. Weakly informative normal priors specified as Normal (0,1/0.01) were used for all intercepts and coefficients. Bayesian inference was implemented using integrated nested Laplace approximation (INLA) (37). This approach uses a deterministic algorithm for Bayesian inference, increasing computational efficiency relative to Markov chain Monte Carlo and other simulation-based approaches (34). We did not explicitly include spatial autocorrelation as several distance-based covariates were included (e.g. distance from own house) (33). Predictive models used data for all individuals aged 8 or over residing in these communities (Table 1) and models were limited to land areas within 5km of households included in the study site. Separate models were fit for each site.

**Table 1.** Baseline characteristics of study site communities and sampled populations

|  | Matunggong |  | Limbuak |  |
| --- | --- | --- | --- | --- |
|  | Sampled | Community* | Sampled | Community* |
| N | 134 | 958 | 109 | 633 |
| Gender |  |  |  |  |
| Male, % (n) | 51.5% (69) | 46.1% (442) | 47.7% (52) | 46.1% (292) |
| Women, % (n) | 48.5% (65) | 53.9% (516) | 52.3% (57) | 53.9% (341) |
| Age in years, median (IQR) | 31 (17 – 53) | 32.5 (8 – 51) | 29 (15 – 46) | 30 (15 – 47) |
| Main occupation, % (n) |  |  |  |  |
| Farming | 29.9% (40) | 28.6% (274) | 7.3% (8) | 10.2% (65) |
| Plantation work | 10.4% (14) | 8.6% (82) | 10.1% (11) | 7.6% (48) |
| Student | 26.1% (35) | 27.7% (265) | 26.6% (29) | 21.0% (133) |
| Other | 6.7% (9) | 9.1% (87) | 15.6% (17) | 14.4% (91) |
| No employment/ housewife | 26.9% (36) | 26.1% (250) | 40.4% (44) | 46.8% (296) |

\* Community includes all individuals eligible for these surveys (residents ages 8 and over)

### Exposure to infected vectors

To estimate vector biting rates, we assembled data from 328 nights of human landing catches (HLCs) conducted with 5km of the Matunggong study site while GPS tracking was on-going, including: monthly longitudinal surveillance (23), investigations surrounding households of cases and controls (24), and environmentally stratified outdoor catches (38) (Supplementary table 2). We limited this data to counts of *An. balabacensis*, the primary *knowlesi* vector which comprises over 95% of *Anopheles* caught in this region. As one experiment only collected mosquitoes for 6 hours, we fit a linear model of all available data vs totals after 6 hour catches to estimate the total numbers of *An. balabacensis* which would have been caught over 12 hours for these data (R^2^= 0.85). Plausible environmental covariates were assembled, including land use type, slope, aspect, elevation, topographic wetness index, EVI, population density and average monthly temperature and rainfall. To select variables for inclusion, Pearson correlation analysis was used to assess multicollinearity between selected environmental variables. As topographic slope and TWI had a strong negative correlation, only TWI was included in the analysis. The autocorrelation function (ACF) and partial autocorrelation function (PACF) were used to explore correlation between time lags.

A Bayesian hierarchical spatiotemporal model was implemented using counts of *An. balabacensis* bites as the outcome, denoted as *m*_*it*_; *j* = 1…n; *t* = 1…n; where *j* indexes location and t indexes month. The log number of person-nights per catch was included as an offset to adjust for numbers of catchers conducting HLCs during different experiments. As the data were over dispersed, a negative binomial distribution was used to model *m*_*it*_. The linear predictor was specified as:

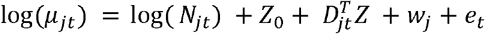

Where *N*_*ijt*_ represents the number of person-nights for each HLC catch, *Z*_0_ represents the intercept, 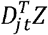 represents a vector of covariates, *w*_*j*_ is the spatial effect and *e*_*t*_ is the temporal effect. The temporal effect e_t_ was included as a fixed effect, random effect or temporally structured random walk model of order 1 in candidate models (39). The spatial effect *w*_*j*_ was modelled as a Matern covariance function between locations *s*_*j*_ and *s*_*k*_:

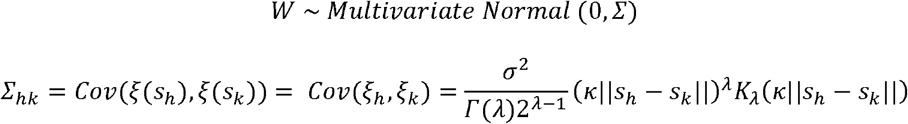

Where ||*s*_*h*_ − *s*_*k*_|| denotes the Euclidean distance between locations *s*_*h*_ and *s*_*k*_, *ξ*(*s*) is the latent Gaussian field accounting for spatial correlation, σ^2^ is the spatial process variance and *K*_*λ*_ is a modified Bessel function of the second kind and order *λ* > 0. κ is a scaling parameter related to *r*, the distance at which spatial correlation becomes negligible, by *r* = √8λ/ κ. A stochastic partial differential equations (SPDE) approach was used, representing the spatial process by Gaussian Markov random fields (GMRF) by partitioning the study area into non-intersecting triangles (40). This approach represents the covariance matrix Σ by the inverse of the precision matrix *Q* of the GMRF (34, 40). Prior distributions were specified on fixed effects and hyperparameters. A vague normal prior distribution was used for the intercept. Weakly informative priors were used for fixed effects specified as *N(1,1/0.01)*. Priors for spatial hyperparameters were specified as range *r* ~ *N(10, 1/0.01)* and standard deviation *σ* ~ *N*(0.1, 1/0.01) as described by Lindgren and Rue (39).

As these vectors are rarely reported indoors (24) and HLCs were primarily conducted outside, we excluded areas within houses for calculations of exposure risks. The proportion of infectious mosquitoes, *c*, was parameterised using a beta distribution for *P. knowlesi* sporozoite rates within this site; with only 4 out of 1524 collected mosquitoes positive, it was not possible to look at variations of infection rates by time and space. Spatially explicit exposure risks were calculated as derived quantity from human resource utilisation, mosquito biting rate models and probability of *P. knowlesi* sporozoite positivity. Individual exposure risk was explored using a simple exposure assessment model where the number of infected bites received by an individual is the sum of bites by infected vector across all locations visited, with the number of infectious bites received by individual *i* in month t as:

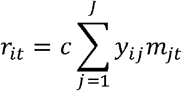

Where *j* indexes the grid cells visited, *y*_*ij*_ is the utilisation distribution, *m*_*jt*_ is the number of bites per individual in that cell and month, and c is the proportion of infectious mosquitoes (4). To evaluate places associated with exposure for the entire community, we calculated the number of infectious bites per grid cell each month as:

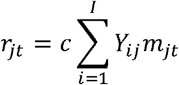

Where *Y*_*ij*_ is the predicted utilisation distribution for all individuals within the community per grid cell *j*. All analyses were conducted in R version 3.5, with Bayesian models implemented using Integrated Nested Laplace Approximation (INLA) (37). Model fit was assessed using deviance information criteria (DIC) and area under the receiver operating curve (AUC), root mean square error (RMSE) or conditional predictive ordinate (CPO) (41).

### Ethics approval

This study was approved by the Medical Research Sub-Committee of the Malaysian Ministry of Health and the Research Ethics Committee of the London School of Hygiene and Tropical Medicine. Written informed consent was obtained from all participants or parents or guardians and assent obtained from children under 18.

## Results

Between February 2014 and May 2016, 285 consenting people participated in the GPS tracking study with 243 included in the final analysis including 109 in Limbuak and 134 in Matunggong (Table 1). The most commonly reported occupation was farm or plantation work (n=73), primarily conducted within the immediate vicinity of the house. A total of 3,424,913 GPS points were collected, representing 6,319,885 person-minutes of sampling time. Median sampling duration was 16.27 days (IQR 13.72 – 19.97), with points recorded for a median of 59.1% (IQR: 46.9% - 71.1%) of the sampling duration. Maximum distances travelled ranged from no travel outside the house to 116km, with a median distance travelled of 1.8km. Utilisation distributions (UDs), the probability of an individual being in a location in space within a given time (Figure 3), varied by gender and occupation. Individuals at the more rural Limbuak site covered larger distances (Table 2), with the largest distances covered by individuals reporting primary occupations of fishing (n=5) and office work (n=9). Although substantial differences were reported in all movements (24 hour sampling) between seasons, no seasonal differences were observed in human movements during peak Anopheles biting times (6pm-6am).

**Table 2.** Home range estimates by demographic group and site

|  | Area of 99% UD for all movement (hectares)<br>Median (IQR) | Area of 99% UD from 6pm – 6am (hectares)<br>Median (IQR) |
| --- | --- | --- |
| Demographic group |  |  |
| Men | 32.09 (7.07, 148.93) | 4.50 (2.79, 19.53) |
| Women | 74.25 (12.24, 320.74) | 6.08 (2.79, 24.17) |
| Children (under 15) | 26.01 (6.39, 151.94) | 3.83 (2.79, 8.73) |
| Occupation |  |  |
| Farming | 29.34 (8.15, 324.38) | 6.75 (2.79, 19.80) |
| Plantation work | 49.14 (9.72, 201.33) | 4.59 (2.79, 27.72) |
| Fishing | 442.49 (40.07, 1189.00) | 227.16 (4.05, 465.14) |
| Office work | 96.80 (63.61, 256.75) | 13.63 (2.88, 20.14) |
| Other | 19.98 (6.30, 26.82) | 2.97 (2.61, 18.27) |
| No employment/ housewife | 43.38 (11.97, 157.59) | 3.60 (2.79, 19.12) |
| Site |  |  |
| Limbuak | 99.99 (24.57, 387.54) | 7.74 (2.88, 58.05) |
| Matunggong | 12.02 (3.94, 85.55) | 2.97 (2.70, 11.77) |
| Season |  |  |
| Dry (February – July) | 28.62 (5.45, 252.45) | 4.19 (2.79, 19.60) |
| Wet (August – January) | 54.90 (17.23, 160.99) | 4.64 (2.79, 19.35) |

**Figure 3.**
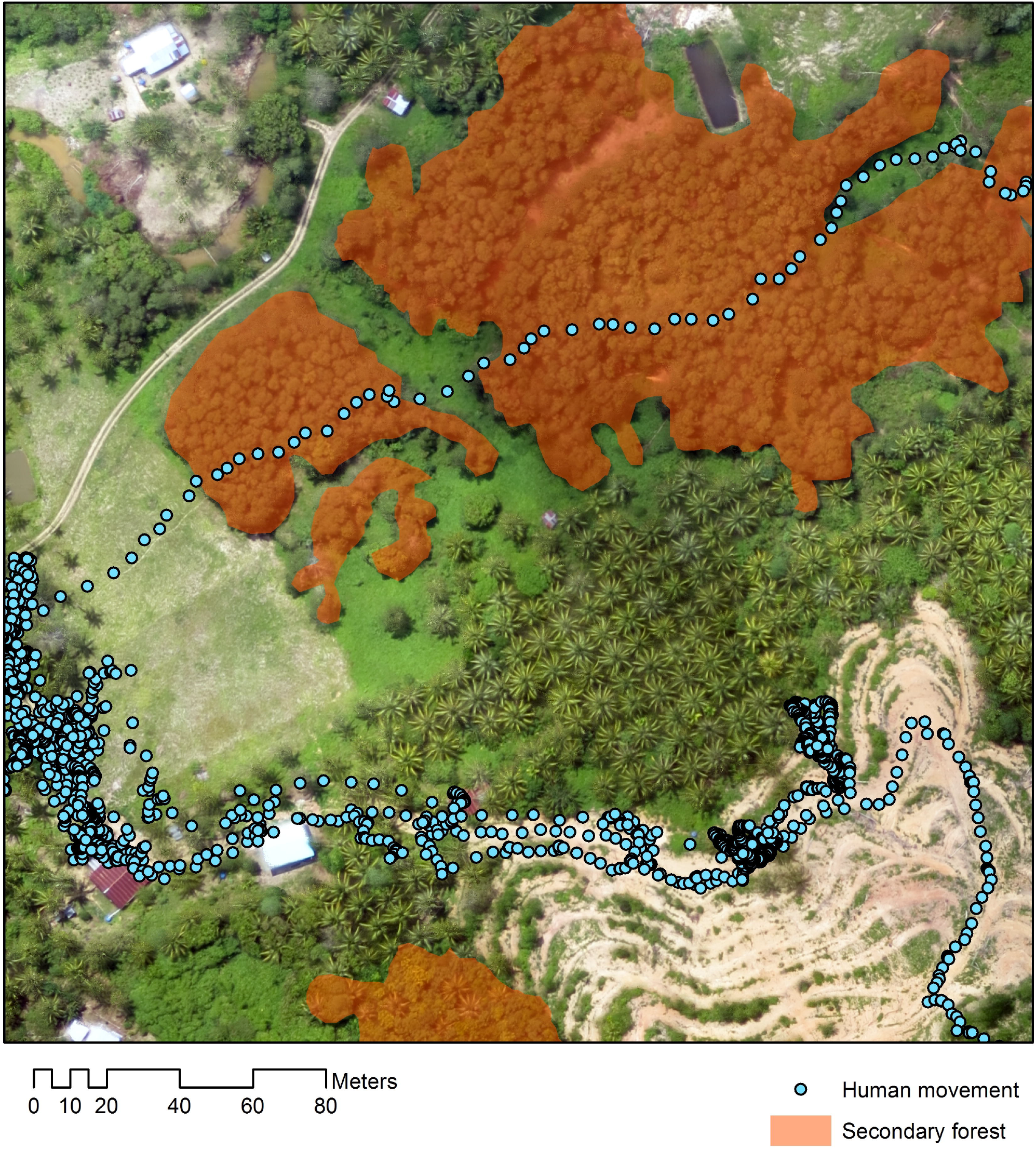
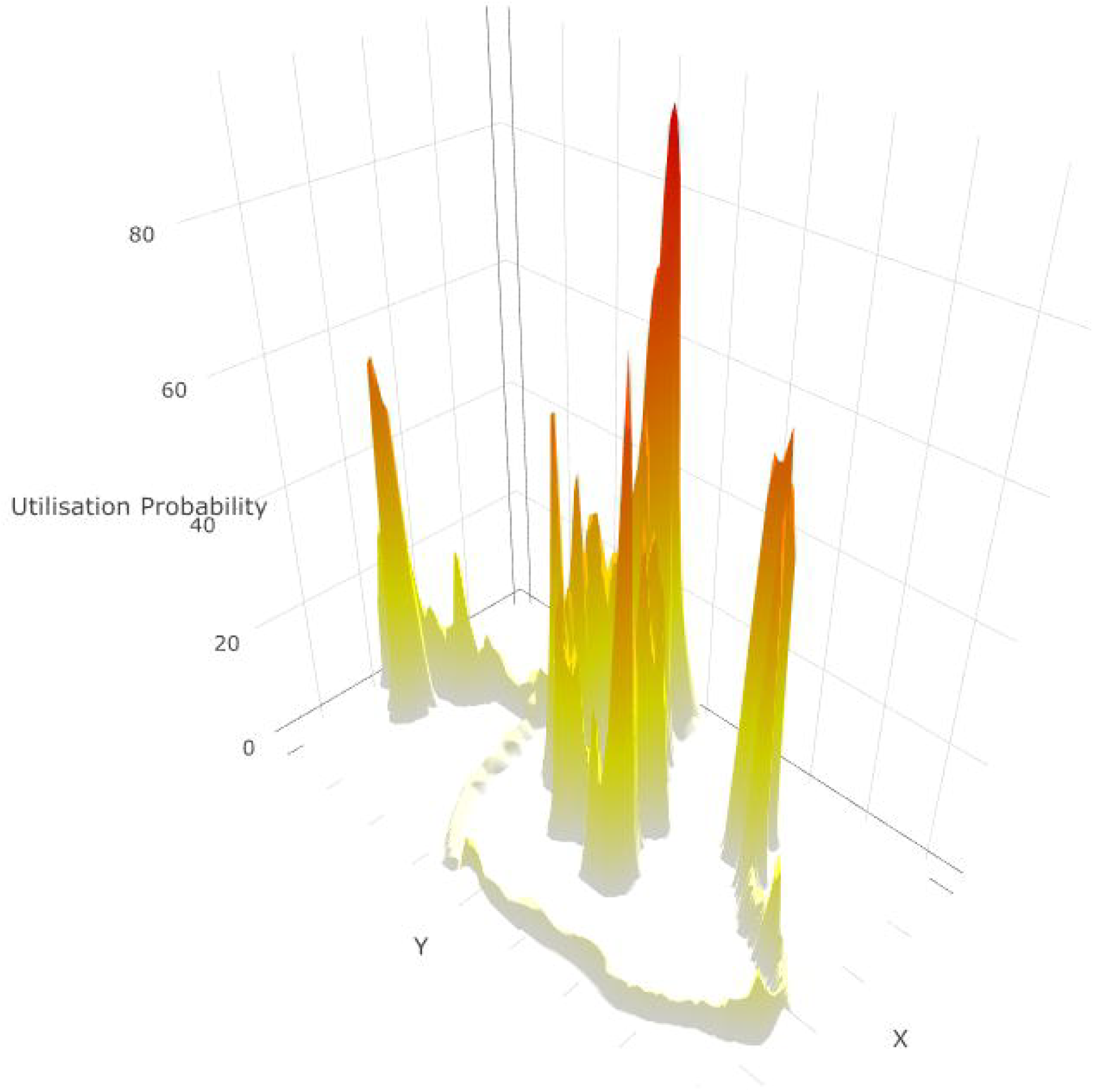
Human movement relative to habitat. A. Example of GPS tracks from a 22-year-old male plantation worker in Matunggong over aerial imagery, B. Individual utilisation distribution calculated from GPS tracks

For both study areas, we developed models of community space use during peak mosquito biting hours (6pm – 6am), in the form of resource utilisation functions, predictions of time- and space-specific UDs on the basis of spatial and environmental variables (28). Between 6pm – 6am, human space use (UDs) was mostly predictable and negatively correlated with distance from the individual’s house, other houses, roads and slope. The AUC for presence/ absence models was 0.936 for Matunggong and 0.938 for Limbuak and RMSE for the overall model was 0.0073 and 0.0043 for Matunggong and Limbuak respectively. While individuals were more likely to use areas further away from forests in the Matunggong site, human space use was positively correlated with proximity to forests in the Limbuak site (Table 3). Despite marked differences between different demographic groups and seasons observed during 24 hour movements, these factors did not improve the predictive power of the model for movements between 6pm and 6am.

**Table 3.** Estimated coefficients for fixed effects of resource utilisation functions (6pm – 6am)

|  | Matunggong |  |  | Limbuak |  |  |
| --- | --- | --- | --- | --- | --- | --- |
|  | Mean | SD | 95% CI | Mean | SD | 95%CI |
| **Probability of presence/ absence** |  |  |  |  |  |  |
| Intercept | 3.383 | 0.839 | 3.218, 3.547 | 3.571 | 0.104 | 3.368, 3.775 |
| Distance from own house (km) | -0.954 | 0.006 | -0.966, -0.942 | -0.543 | 0.003 | -0.548, -0.539 |
| Distance from forest (km) | 5.997 | 0.177 | -5.650, 6.344 | -1.845 | 0.050 | -1.944, -1.746 |
| Distance from road (km) | -5.552 | 0.057 | -5.663, -5.441 | -3.656 | 0.019 | -3.694, -3.618 |
| Distance from houses (km) | -0.504 | 0.030 | -0.563, -0.444 | 0.176 | 0.007 | 0.162, 0.189 |
| Elevation (100 MSL) | -0.710 | 0.025 | -0.759, -0.662 | -1.268 | 0.037 | -1.340, -1.197 |
| Slope (degrees) | -0.0244 | 0.002 | -0.028, -0.021 | -0.009 | 0.001 | -0.012, -0.006 |
| **Utilisation distributions for locations present** |  |  |  |  |  |  |
| Intercept | -6.846 | 0.866 | -8.549, -5.147 | -5.676 | 1.017 | -7.673, -3.681 |
| Distance from own house (km) | -0.583 | 0.004 | -0.590, -0.576 | -0.308 | 0.002 | -0.311, -0.305 |
| Distance from forest (km) | 12.012 | 0.199 | 11.621, 12.403 | -1.771 | 0.049 | -1.868, -1.675 |
| Distance from road (km) | -0.833 | 0.054 | -0.939, -0.728 | -1.532 | 0.011 | -1.554, -1.511 |
| Distance from houses (km) | -0.819 | 0.023 | -0.864, -0.773 | -0.239 | 0.006 | -0.249, -0.228 |
| Elevation (100 MSL) | 0.664 | 0.027 | 0.610, 0.718 | -0.297 | 0.003 | -0.303, -0.297 |
| Slope (degrees) | -0.021 | 0.002 | -0.024, -0.018 | -0.034 | 0.001 | -0.036, -0.031 |

Between August 2013 and December 2015, 4814 *An. balabacensis* were caught from 328 sampling nights in 155 unique locations. The median biting rate was 2.1 bites per night per person, ranging from 0 – 28 bites per person per night (Figure 4). Despite monthly variation, no significant time lags or seasonal patterns in mosquito biting rates were identified (Table 4). Although no associations were identified between land classification and vector density in this site, models identified positive relationships with enhanced vegetation indices (EVI) and negative associations with distance to forest and human population density (Table 5). Of 1524 mosquitoes tested for *Plasmodium* sporozoites, the median sporozoite rate was 0.24% (95% CI: 0.09 – 0.58%).

**Table 4.**
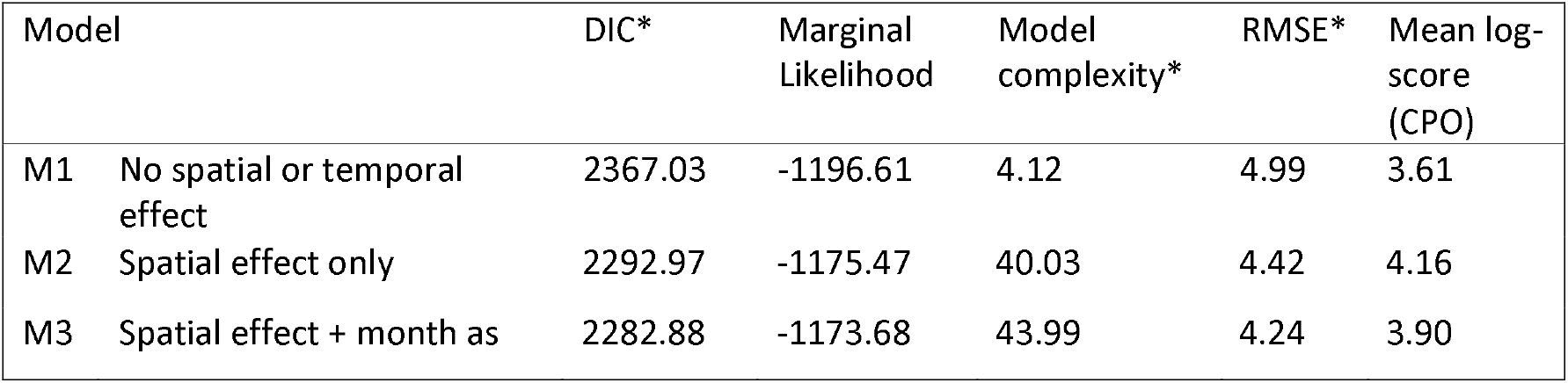
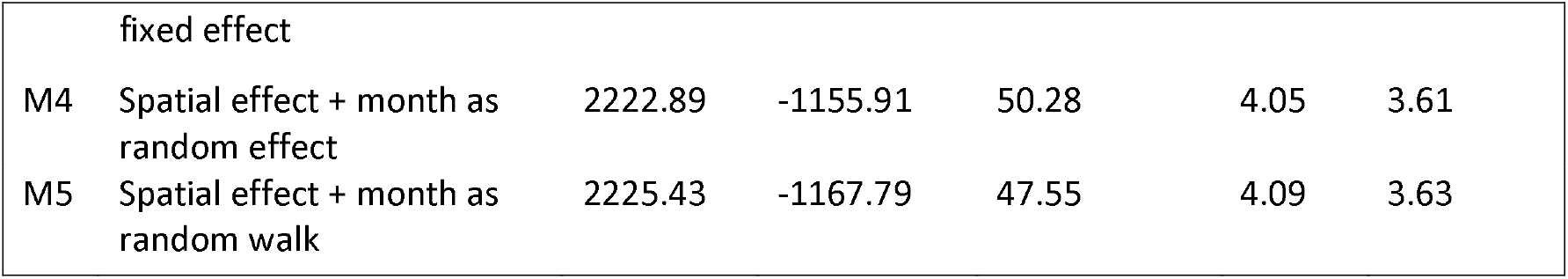
Model selection statistics for mosquito biting rates

| Model | DIC* | Marginal Likelihood | Model complexity* | RMSE* | Mean log-score (CPO) |
| --- | --- | --- | --- | --- | --- |
| M1 No spatial or temporal effect | 2367.03 | -1196.61 | 4.12 | 4.99 | 3.61 |
| M2 Spatial effect only | 2292.97 | -1175.47 | 40.03 | 4.42 | 4.16 |
| M3 Spatial effect + month as | 2282.88 | -1173.68 | 43.99 | 4.24 | 3.90 |

|  |  |  |  |  |  |  |
| --- | --- | --- | --- | --- | --- | --- |
|  | fixed effect |  |  |  |  |  |
| M4 | Spatial effect + month as random effect | 2222.89 | -1155.91 | 50.28 | 4.05 | 3.61 |
| M5 | Spatial effect + month as random walk | 2225.43 | -1167.79 | 47.55 | 4.09 | 3.63 |

**Table 5.** Posterior rate ratio estimates and 95% Bayesian credible interval (BCI) for model 4 of mosquito biting rates

| Covariate | 95% BCI Rate Ratio |  |  |
| --- | --- | --- | --- |
|  | Mean | 2.5% | 97.5% |
| Population density | 0.963 | 0.916 | 1.004 |
| EVI | 3.185 | 1.185 | 8.532 |
| Distance to forest (100m) | 0.926 | 0.871 | 0.976 |
| Spatial range (km) | 3.120 | 0.514 | 6.926 |

**Figure 4.**
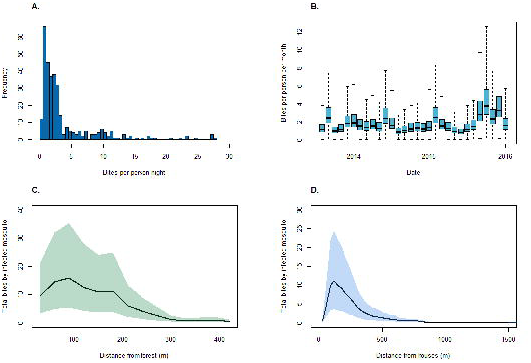
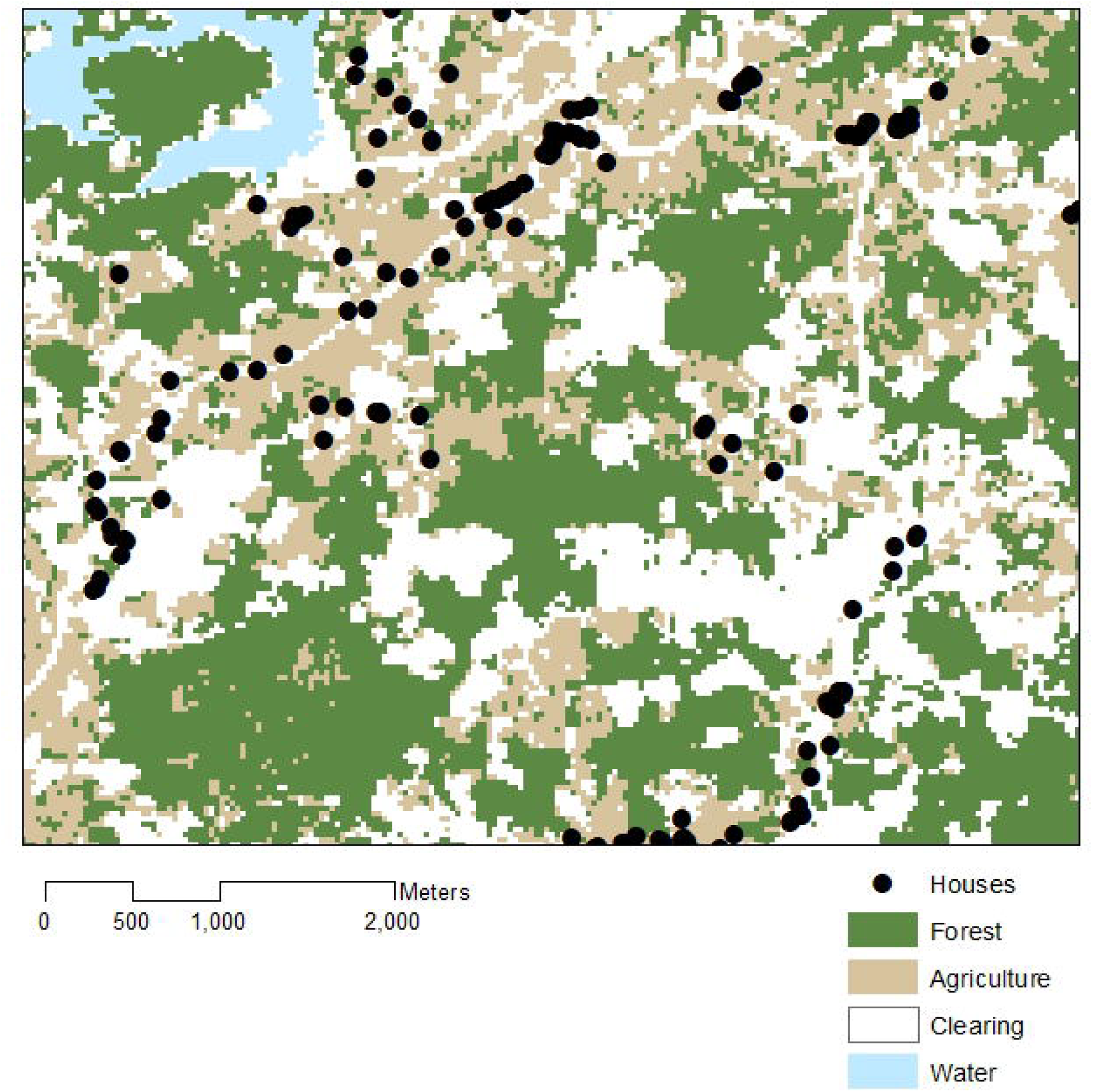
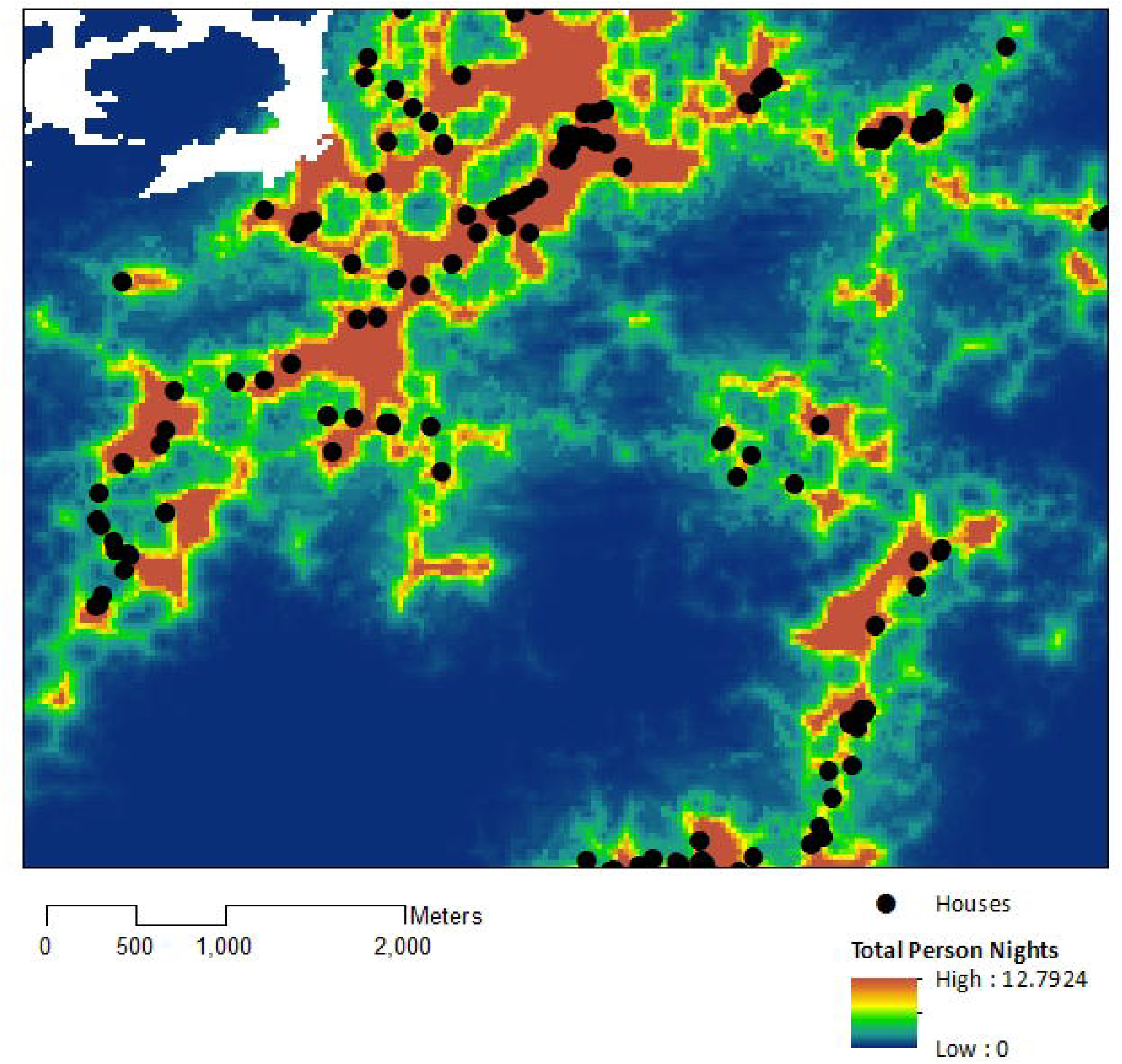
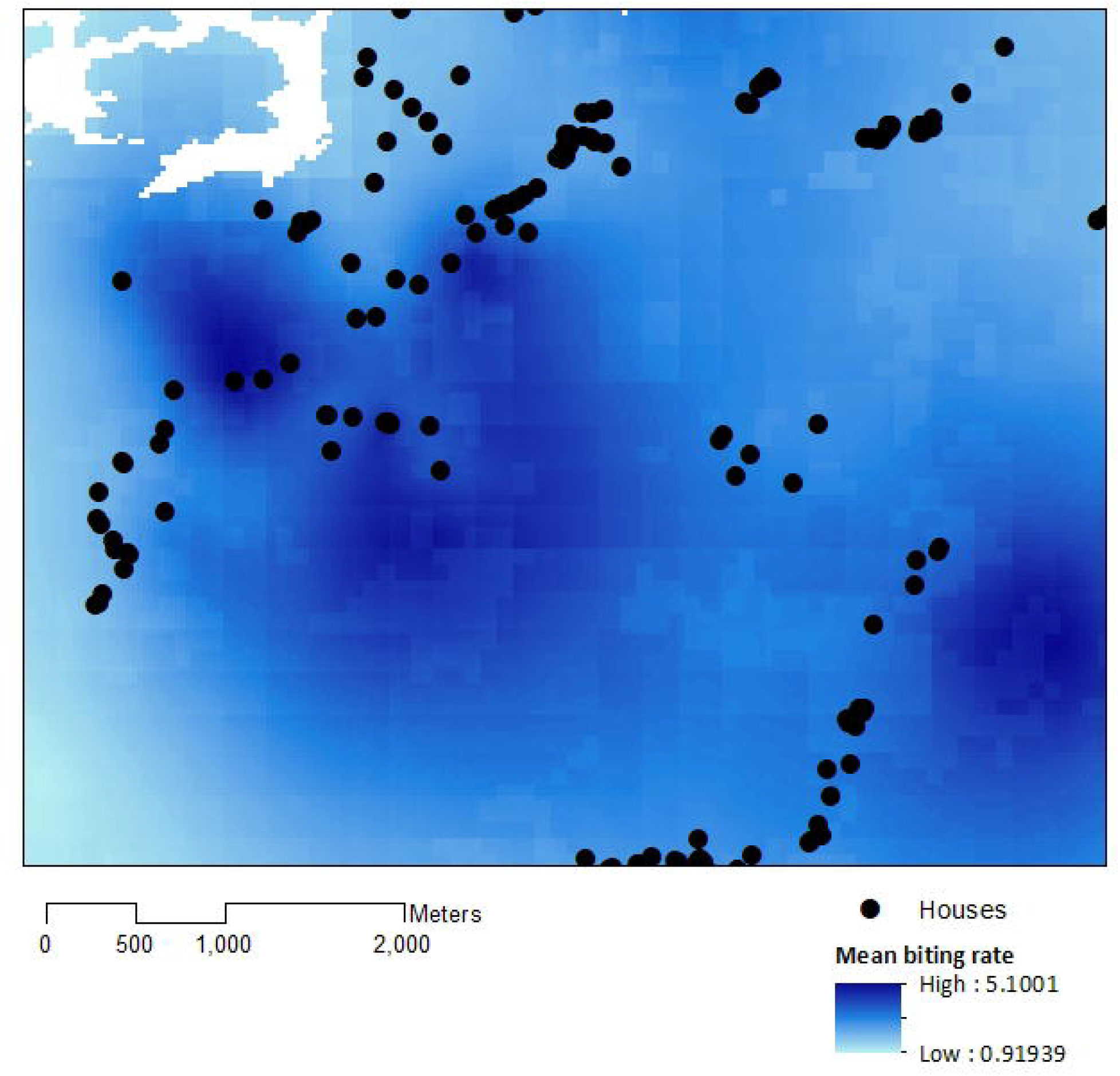
Mosquito biting rates. A. *An. balabacensis* biting rate per person-night from data collected in Matunggong, B. Predicted mean *An. balabacensis* biting rates per month from spatiotemporal models, C. Predicted number of bites for all individuals residing in Matunggong by distance from secondary forest, and by D. Distance from households

For individuals included in the GPS tracking study in Matunggong, where both human movement and entomology data was available, we calculated exposure risks as a derived quantity from utilisation distributions and mosquito biting rate models. Exposure varied markedly between individuals, with an overall 150-fold difference in predicted mean probabilities of infected bites per night (range: 0.00005-0.0078) (Table 6). No clear differences were observed between genders, age groups or occupations of individuals sampled and there was no association between risk and distance travelled.

**Table 6.** Probabilities of infected bites per night for sampled individuals in Matunggong by demographic characteristics

|  | Predicted infectious bites per night (median (IQR)) |
| --- | --- |
| Demographic group |  |
| Men | 0.00157 (0.000804, 0.00289) |
| Women | 0.00219 (0.000864, 0.00307) |
| Children (under 15) | 0.00131 (0.000812, 0.00330) |
| Occupation |  |
| Farming | 0.00180 (0.00101, 0.00362) |
| Plantation work | 0.00216 (0.000680, 0.00278) |
| Student | 0.00143 (0.000915, 0.00304) |
| Other | 0.00225 (0.000852, 0.00302) |
| No employment/ housewife | 0.00142 (0.000297, 0.00263) |

Using the resource utilisation function with demographic and spatial data for all individuals in Matunggong, we predicted community-wide space use and estimated exposure to infected mosquitoes (Figure 5). The predicted number of person nights per grid cell for the entire community ranged from 0 to 12.79 (median: 0.01, IQR: 0.0004 – 0.99), with the mean probability of a community member exposed to an infected bite per grid cell of 0.00082 (IQR: 0.00001, 0.00050). Although over 43% of the study site is forest and relatively high biting rates were predicted in forests during the study period (mean: 1.94, range: 0.04 – 12.59), this habitat was rarely used by people in the evenings, with less than 8% of predicted person-nights in forests. Models only based on mosquito biting rates and not including human space use predicted 42% of infectious bites occurred in forested areas and only 8.6% of bites occurring within 100m of houses (Figure 5C). In contrast, when space use patterns are included, over 91% of predicted infected bites were predicted within 500m of houses (Figure 5D). Highest exposure risks were consistently found near forest edges and in close proximity to households, despite spatial and temporal heterogeneity and model uncertainty (Figure 4).

**Figure 5.**
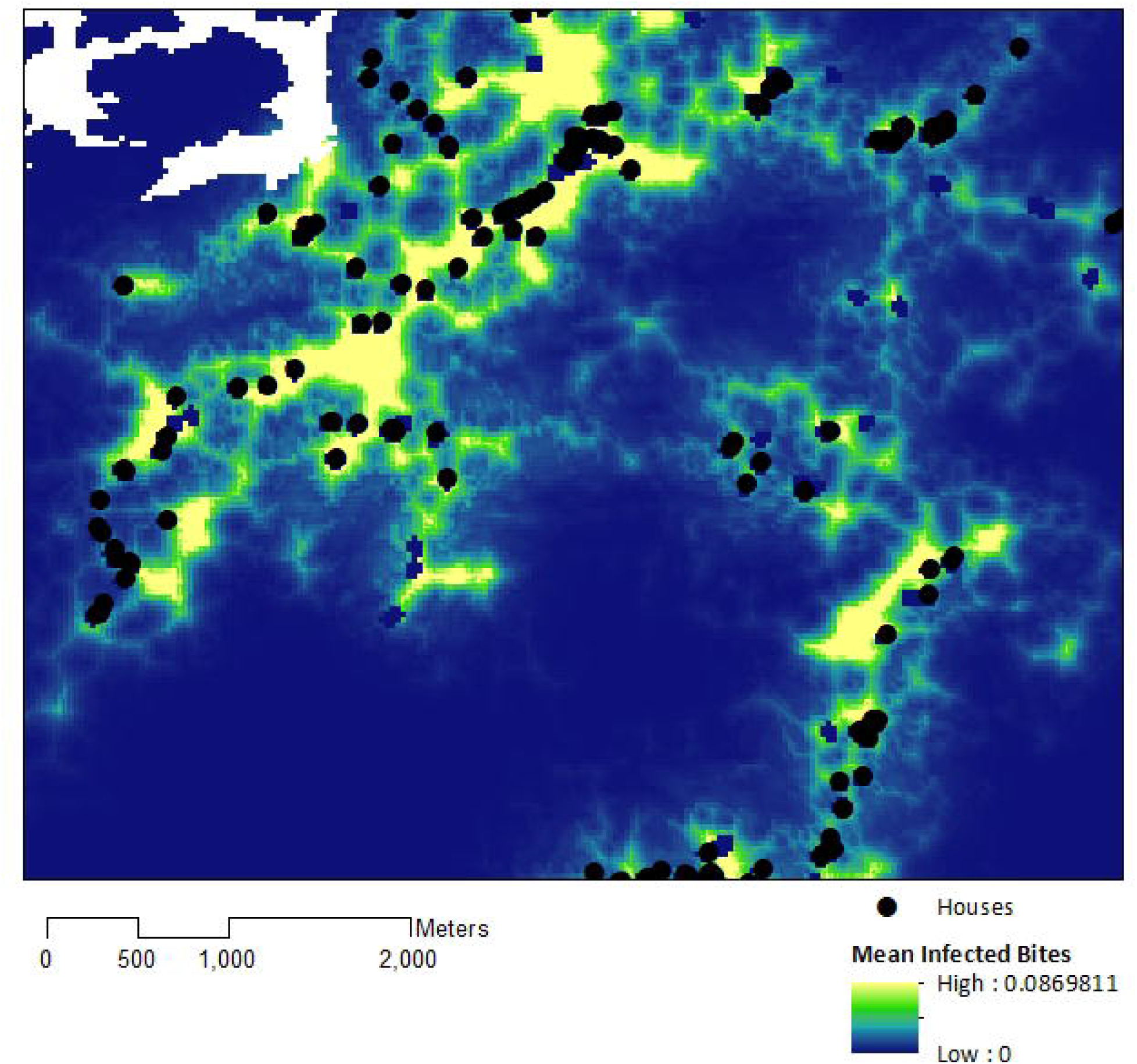
A. Land use in Matunggong site, B. Predicted number of person-nights for entire community per grid cell, C. Predicted mosquito biting rates, D. Predicted infected bites per grid cell

## Discussion

This study highlights the importance of human space use in different land cover types in determining exposure to zoonotic and vector-borne diseases such as *P. knowlesi*. Although *P. knowlesi* has previously been associated with forest exposure (e.g. (19)) and higher biting rates have been reported in forest interiors (23), this novel approach incorporating both mosquito and human space use data provides a new perspective on peri-domestic transmission, with more than 90% of infectious bites predicted in areas surrounding households at forest edges. This study additionally demonstrates the utility of ecological methods to understand human movement and identify geographical areas associated with higher contact with disease vectors.

Within these communities, local movement patterns during peak vector times were largely predictable and could be explained by spatial and environmental factors. However, despite this finding, there was substantial variation in predicted exposure between individuals as a result of heterogeneity in habitats used. No significant differences in exposure were predicted between men and women, with individuals with high exposure risks identified across occupational and age groups. Although this finding differs from clinical reports, a comprehensive survey within this community identified equal proportions of men and women exposed to *P. knowlesi* as evidenced by specific antibody responses and data on asymptomatic infections suggests higher numbers of non-clinical infections in women (30, 42). While infrequent events or long-range movements (such as hunting trips) may contribute to these differences in clinical cases and may not have been captured within this two-week sampling period within the study site, this analysis highlights the importance of routine movements into local environments in shaping exposure risks.

This improved understanding of how local human land use is related to exposure risk has important implications for surveillance and control programmes. Malaria control programmes often rely on interventions within the house, such as insecticide treated bednets and indoor residual spraying; however, movements outside during peak biting times illustrate the importance of also targeting outdoor transmission. The identification of areas where exposure is likely to occur can further be used to refine interventions; for example, although insecticide treated hammocks have been proposed for deep forest environments, larval source management may be more appropriate to target environments in close proximity to houses. Although initial *P. knowlesi* cases were primarily identified in adult men living and working in forests (20), this study illustrates the potential importance of peri-domestic habitats in transmission and provides quantitative insight on mixing between people and infected mosquitoes in forest fringe areas. As Malaysia moves towards malaria elimination, surveillance systems are incorporating novel focal investigation methods, including monitoring changes in local land use and populations at risk (43). In additional to routine vector surveillance, this study highlights the need to incorporate measures of human space when defining risk zones.

Even with the large and highly detailed movement dataset analysed, this study was limited by the availability of mosquito data; as human landing catch data were assembled from other studies, there was not uniform spatial and temporal coverage of the study site increasing uncertainty. The limited mosquito data availability precluded development of mosquito biting rate models for Limbuak and other outlying islands. An additional limitation to estimating mosquito biting rates was the difficulty obtaining spatially and temporally resolute remote sensing data for predictors due to high cloud cover (44). As few positive mosquitoes were identified, uniform estimates of sporozoite rates based on available data were used across the Matunggong site; if further data was available, these models could be refined to incorporate estimates of human and macaque density, mosquito biting preferences in different habitats and infection levels in all hosts (45). Additionally, as this study was designed to quantitatively estimate time spent in different landscapes, further studies could explore other aspects of land use, such as the purposes of travel, activities undertaken or practices used to modify or management land cover.

Despite these limitations, this is the first large-scale study to utilise GPS tracking data and ecological methods to create fine-scale maps of exposure risk. This study highlights the importance of incorporating heterogenous patterns of human space use into disease models, as the majority of human exposure may occur in areas with lower vector biting rates but greater probabilities of human use. Further, results quantitatively illustrate the importance of forest edges and local habitat in *P. knowlesi* transmission and can inform understanding of other zoonotic and vector-borne diseases.

## Acknowledgements

We would like to thank the MONKEYBAR project team and the participants in Sabah, Malaysia for their help with these studies. We acknowledge the Medical Research Council, Natural Environmental Research Council, Economic and Social Research Council and Biotechnology and Biosciences Research Council for funding received for this project through the Environmental and Social Ecology of Human Infectious Diseases Initiative, grant no. G1100796. This work was additionally supported by the National Socio-Environmental Synthesis Center (SESYNC).

