## Supplementary File for "Local human movement patterns and land use impact exposure to zoonotic malaria in Malaysian Borneo"

**Supplementary file 1A. Data sources for assessed spatial and environmental covariates**

| Covariate | Description | Spatial resolution | Source | Reference |
|---|---|---|---|---|
| Land use type | Classified land use type as forest, agriculture, clearing and water | 30m | Derived from Landsat, described in Fornace et. al, *Plos Neglected Tropical Diseases*, 2018 | Landsat 8 Operational Land Imager. NASA EOSDIS Land Processes DAAC, USGS Earth Resources Observation and Science (EROS) Center. 2014. Available from: http://landsat.usgs.gov//index.php. |
| EVI | Monthly mean enhanced vegetation index (0-1) | 250m | MODIS Vegetation Indices | DAAC NL. MODIS/ Terra Vegetation Indices 16-Day L3 Global 250m Grid SIN V006. Sioux Falls, South Dakota: USGS Earth Resources Observation and Science (EROS) Center. |
| TWI | Topographic wetness index | 30m | Calculated from ASTER GDEM | Advanced Spaceborne Thermal Emission and Reflection Radiometer Global Digital Elevation Model (ASTER GDEM) Version 2 [Internet]. NASA EOSDIS Land Processes DAAC, USGS Earth Resources Observatoin and Science (EROS) Center. 2015. Available from: http://gdem.ersdac.jspacesystems.or.jp/ |
| Elevation | Meters above sea level (m) | 30m | ASTER GDEM | |
| Slope | Degrees of slope incline (degree) | 30m | Calculated from ASTER GDEM | |
| Aspect | Compass direction slope is facing (degree) | 30m | Calculated from ASTER GDEM | |
| Distance to houses | Euclidean distance from nearest household, calculated from household GPS points (m) | 1m | Mapped during study using Garmin GPS, calculated in R | |
| Distance to roads | Euclidean distance from nearest road, calculated from GPS tracks of roads (m) | 1m | Mapped during study using Garmin GPS, calculated in R | |
| Population density | Estimated population per 100m for 2015 (adjusted for UN estimates) | 100m | WorldPop | Lloyd CT, Sorichetta A, Tatem AJ. High resolution global gridded data for use in population studies. Sci Data. 2017;4:170001. |
| Precipitation | Average monthly rainfall (mm/ day) | 0.25° | Tropical Rainfall Measuring Mission | Tropical Rainfall Measurement Mission Project (TRMM). Daily TRMM and other satellites precipitation product (3B42 V6 derived). In: DAAC) GSFCDAACG, editor. 2015. |
| Temperature | Average monthly temperature (°C) | - | Malaysian Meteorology Department | |

**Supplementary file 1B. Data sources of mosquito biting data**

| Study | Study design | Sampling dates | Collection time | Data points | References |
|---|---|---|---|---|---|
| Longitudinal sampling | Longitudinal monthly sampling at sentinel sites in Matunggong and Limbuak | August 2013-December 2014 | 12 hours, 6pm – 6am | 82 | Chua TH, et al. Phylogenetic analysis of simian Plasmodium spp. infecting Anopheles balabacensis Baisas in Sabah, Malaysia. *PLoS Neg Trop Dis*. 2017;11(10):e0005991.<br><br>Wong ML, et al. Seasonal and Spatial Dynamics of the Primary Vector of Plasmodium knowlesi within a Major Transmission Focus in Sabah, Malaysia. *PLoS Neg Trop Dis*. 2015; 9(10):e0004135. |
| Case control | Sampling around houses of *P. knowlesi* cases and matched controls | February 2014 – July 2014 | 12 hours, 6pm – 6am | 34 | Manin BO, Ferguson HM, Vythilingam I, Fornace K, William T, Torr SJ, et al. Investigating the Contribution of Peri-domestic Transmission to Risk of Zoonotic Malaria Infection in Humans. *PLoS Neg Trop Dis*. 2016;10(10):e0005064. |
| Environmentally stratified sampling | Monthly sampling in randomly selected 100m$^2$ grid cells, stratified by land cover | February 2015 – December 2015 | 6 hours, 6pm – 12am | 212 | Ng SH, Homathevi R, Chua TH. Mosquitoes of Kudat: species composition and their medical importance (Diptera: Culicidae). *Serangga*. 2016;21(2):149-62. |
